# Brain dynamics during architectural experience: prefrontal and hippocampal regions track aesthetics and spatial complexity

**DOI:** 10.1101/2025.01.09.631831

**Authors:** Lara Gregorians, Zita Patai, Pablo Fernandez Velasco, Fiona E. Zisch, Hugo J. Spiers

## Abstract

Architectural experience involves processing the spatial layout of an environment and our emotional reaction to it. However, these two processes are largely studied separately. Here we used functional magnetic resonance imaging (fMRI) and first-person movies of journeys through buildings and cities to determine the contribution of different brain regions to spatial and aesthetic aspects of the built environment. During scanning, participants watched 48 movies that show first-person-view travel through different spaces; immediately after each video, they either judged the spatial layout complexity or valence of the environment. After scanning, participants also reported the memorability of the spaces encountered. Activity in brain regions previously linked to valence processing (e.g. ventromedial prefrontal cortex) were modulated by aesthetic qualities of the stimuli (i.e. increased for pleasant spaces compared to unpleasant spaces) and the task (more active when judging valence), whereas activity in brain regions linked with spatial processing (e.g. parahippocampal regions) increased in complex layouts compared to simple layouts. The hippocampus and parahippocampal cortex were associated with the memorability of spaces and were modulated by both aesthetic and spatial qualities. We also tested for curvature, fascination, coherence and hominess - qualities linked to aesthetic judgement in architecture. We replicated findings activating right lingual gyrus for fascination, left inferior occipital gyrus for coherence, and left cuneus for hominess, and found inverse curvature (increasing rectilinearity) activated spatial, valence and visual processing regions. Overall, these findings provide important insights into how different brain regions respond whilst experiencing new buildings and city spaces, which is needed to advance the field of neuroarchitecture.

## Introduction

Moving through the built environment involves entwining many processes simultaneously – cognitive, affective and behavioural^1–4^. These processes span both the coding of relevant spatial properties of the environment and one’s affective engagement with its aesthetics properties. And yet, how the built environment gives rise to a variety of brain responses remains poorly understood. Many experiments have investigated how our brains code spatial information in our surroundings^5^ and some studies have examined how our brains process the aesthetic and affective qualities of architectural images^2^. So far, however, these two aspects – spatial and affective processing – have not been investigated together. Whilst the neural correlates of aesthetic and affective experience are often considered in relation to one another, no studies to date have examined how they relate to the neural correlates of spatial cognition. Affect considers facets of emotions, and aesthetic experience relates to the experience of beauty and pleasure. This can be formalised by studies asking about the beauty of a stimulus (aesthetic experience) or like/dislike towards a stimulus (‘valence’, affective experience). Moreover, studies exploring affect and aesthetic experience often use still images, which lack the dynamic nature of real-world experiences of moving through architectural spaces^4^. While static images can provide valid ecological stimuli for other forms of aesthetic experience, such as paintings, architectural experience is intrinsically dynamic. We rarely just look at buildings - we experience them by moving in and through them^4^. Therefore, this study builds off of our previous work in which we sourced, curated and evaluated a database of videos that show first-person journeys through built environments, which vary in valence and spatial complexity^4^. Here, we used these videos of journeys through buildings to explore the brain regions involved in both aesthetic and affective, as well as spatial processing of the built environment.

A substantive body of work in neuroscience has examined the brain structures involved in spatial mapping and navigation. For novel or recently learnt environments, navigation is heavily reliant on the hippocampus^6–8^, which is thought to encode a ‘cognitive map’ representing the structure of the environment^9^. The hippocampus operates along with a broader circuit of brain regions for spatial mapping and navigation, including the parahippocampus, the retrosplenial cortex (RSC), and the occipital place area (OPA)^5,8,10–15^. The parahippocampus has been linked to scene and landmark processing, and the RSC integrates this scene within the cognitive map to aid navigation^16–25^. As for the OPA, this region is key in mapping the navigational affordances of real-world spaces – e.g. available paths and boundaries – and is involved in scene processing together with the parahippocampal place area (PPA) and RSC^3,25–28^. The above studies have paid little attention to affect, with only a few experiments considering how emotional input impacts our memories of space^29–31^.

Aesthetic qualities are central to affect in the built environment^2^, whether it is in the engagement with aesthetic atmospheres^32^, meaning making^33^ or our movement through space^34^. Valence, or pleasantness, is central to aesthetic processing^35^, and is the most widely studied dimension of affect^36^. The core circuit for aesthetic and valence processing consists of the ventromedial prefrontal cortex (vmPFC), the orbitofrontal cortex (OFC), the anterior cingulate cortex (ACC), the amygdala, and the insula^35,37–39^. The vmPFC is associated with value judgments, reward-related decision making, and positive valence^40–43^. It receives inputs from the OFC, which appears to track valence across different stimuli^44–46^. Both the OFC and vmPFC connect directly to the ACC, further supporting reward processing^46^. The amygdala is associated with memory retrieval of emotionally enhanced events^47^ and has classically been tied to fear-encoding and to the processing of negative valence or emotion^48^. More recently, it has been shown to also respond to positively valenced stimuli^49,50^. Likewise, the insula has traditionally been associated with negative feelings, but it has now been found to correlate with the processing of both positively and negatively valenced stimuli^35,51^.

Alongside the core circuit of aesthetic processing, there is some evidence that points to the involvement of brain regions typically involved in spatial mapping in valence processing too. The hippocampus, parahippocampus and RSC have all previously been found to process the emotional significance of words^52–54^. The hippocampus has been linked to encoding valenced, non-arousing information^55^. Parahippocampal activity responds to viewing unpleasant pictures, increasing arousal for pleasant pictures, and emotive words^56–58^. It may be modulated by valence, activating in relation to approach judgments for positive social scenes compared to negative social scenes in approach-avoidance tasks^59^. Previous studies have also found parahippocampal activity when participants viewed pictures of rooms which had been previously associated with negatively arousing events^29^. Nonetheless, there does not seem to be enough evidence yet to suggest that these regions - traditionally associated with scene processing and spatial memory - are central to aesthetic processing across domains.

A notable recent development in the study of architectural experience comes from the work of Coburn and colleagues^60^, who found that aesthetic responses to images of architectural interiors can be largely explained through three psychological dimensions: fascination, coherence and hominess. Fascination corresponds to the interest that the environment generates, coherence is the ease with which a participant comprehends an environment, and hominess is the degree to which the environment feels like a personal space. These three dimensions all correlated with valence, but evoked distinct neural responses depending on whether the task at hand was aesthetic evaluation, or approach-avoidance judgements.

Coburn and colleagues also explored how salient design features, such as curvature, modulated these three dimensions of architectural experience^60^. Curvature is one of the key physical features studied in neuroarchitecture and architectural psychology^39,60–66^ given that contour is an important component of the composition of space, and has also been studied extensively in different applications of neuroaesthetics^39,63,67–69^. Curvature can be calculated computationally by extracting the contours of an image or can be deciphered by expert judgments (e.g. architects categorising images of spaces as curvilinear or rectilinear). Both metrics of curvature have been found to correlate with aesthetic responses in architecture^39,64–66^ as well as other domains (e.g. object shapes)^63,67–69^. Typically curved spaces are considered more beautiful and pleasing^39,65,66^. Therefore, in a further effort to bridge our understanding of affective, aesthetic and spatial processing, we considered fascination, coherence, hominess and curvature as key factors that relate to aesthetic judgments in spaces. We note that there are many other architectural features that could also be studied alongside curvature (e.g. scale, lines of sight), but given the scope of this work, we explore one feature that has empirically driven hypotheses for neural correlates.

In the present work, participants were scanned with functional magnetic resonance imaging (fMRI) while they watched videos of first-person trajectories through built environments and rated either the valence or the spatial layout complexity of each space (figure 1). We employed a 2 x 2 x 2 design, where movies were either positive or negative in valence (based on prior ratings^4^), complex or simple (based on prior ratings^4^) and movies were either judged on valence or complexity (the task manipulation). Participants were split into two participant groups (PG1 and PG2) with inverse task foci for counterbalancing (e.g. movie 1 was judged for valence by PG1, and for spatial complexity by PG2). Past ratings of the movies were used to derive measures of hominess, coherence and facination^4^, and video curvature was calculated from image contours (see figure 1B for examples of variation in curvature). The primary aim of this study was to explore the brain regions that support affect and spatial mapping, and determine how these networks operate depending on the affect-value of the space and on the task that participants are asked to focus on. Following a post-scan debrief session, participants also judged which spaces were most memorable to help understand the brain regions involved in encoding memories of spaces, and to explore the qualities of spaces that impact this.

**Figure 1.**
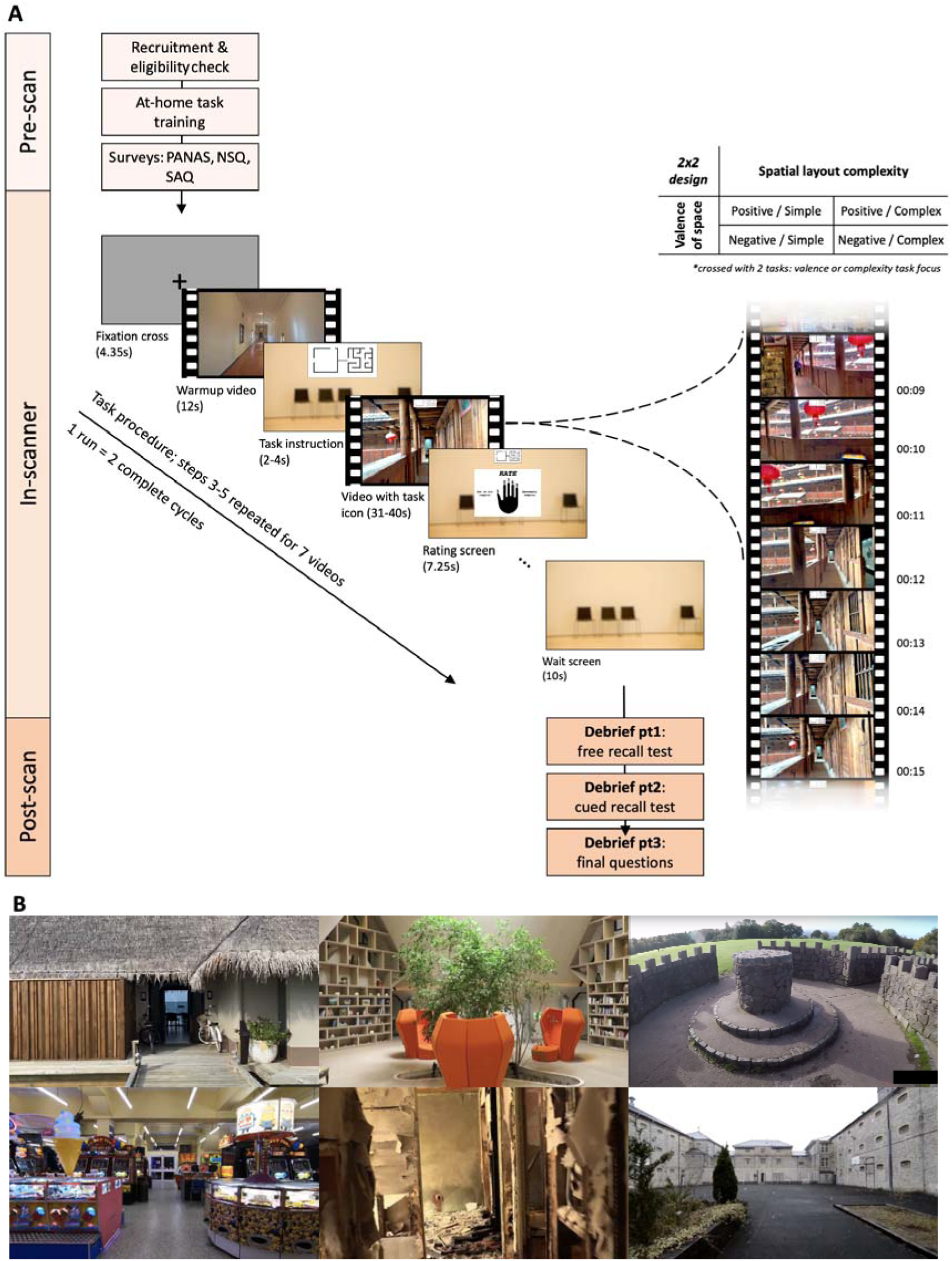
Experimental Task. (A) Task structure. In-scanner the participant saw a fixation cross followed by a warm-up video of progress down a corridor. They then saw the task instruction screen, indicating whether they should focus on valence or spatial complexity in the upcoming video. The video then played. At the end, the participant was asked to give a valence or complexity rating for that video (1 = low, 5 = high). This cycle repeated for seven videos, after which they got a short break while a waiting screen displayed. This instruction-video-rating process then repeated with seven new videos. Two complete cycles equated to one run; one run had 14 videos excluding warm-up; four runs total. Participants were split into two participant groups (PG1 and PG2) with inverse task foci - i.e. if PG1 rated video 1 for valence, PG2 rated video 1 for spatial complexity. PANAS = Positive and Negative Affect Schedule^72^. NSQ = Navigational Strategy Questionnaire^73^. SAQ = Spatial Anxiety Questionnaire^74^. **(B) Stills from 6 example videos.** 48 videos were selected from our database of valenced videos. Stills represent spaces with high and low valence and spatial complexity. Images in the third column demonstrate spaces with high and low curvature.

Building on prior research, we hypothesised that brain networks associated with spatial mapping (hippocampus, parahippocampal gyrus, OPA and RSC) would respond to the spatial layout complexity of architectural spaces and that these networks would respond more strongly when participants were asked to focus on the spatial layout complexity of the spaces. Similarly, we hypothesised that brain networks associated to valence and reward processing (ACC, OFC, vmPFC, amygdala and insula) would respond to the pleasantness of architectural spaces and that they would respond more strongly when participants were asked to focus on the pleasantness of the spaces. Following prior work suggesting distinct activation of anterior temporal and posterior medial processing networks to memory^70^, we considered that there may be different responses in anterior and posterior regions in relation to aesthetic and spatial mapping. We also investigated whether ratings of fascination, coherence, hominess and metrics of curvature - key factors of aesthetic experience in architecture^60^ - would show modulation in the brain networks noted for aesthetics and spatial processing, as well as regions previously associated with these factors. These are the right lingual gyrus for fascination, left inferior occipital gyrus for coherence, and left cuneus for hominess^60^. And, given past research has reported curvature to activate the visual cortex (for approach-avoidance decisions and beauty judgments^61^) and the ACC (for beauty judgments^39^), we explored whether we would observe increased responses to curvature in these regions. We ran small volume corrections on regions-of-interest (ROI) for all hypothesised brain regions and whole-brain family-wise error (FWE) correction (p<.05, minimum voxel cluster threshold >5) for visual cortex. Finally, we also examined what makes some places more memorable, as well as what potential correlations exist between memorability and aesthetic, affective and spatial metrics derived for each video^4^. Prior work suggests that movies with high affective qualities and distinctiveness will be better remembered^30,71^.

## Results

### Brain imaging results

We first present our fMRI results and then turn to discoveries from participant behaviour. Our valence-related ROIs were the ACC, OFC, vmPFC, insula and amygdala, and our spatial processing ROIs were the hippocampus, OPA, parahippocampal gyrus and RSC. In summary, we found activation for valence (vs spatial) task focus for all of our valence ROIs; activation for positive (vs negative) spaces in the vmPFC and in all of our spatial ROIs; activation for complex (vs simple) spaces in the hippocampus and parahippocampus; activation for inverse curvature (increasing rectilinearity, see figure 1B for example spaces) in the amygdala, insula and vmPFC, as well as in hippocampus, parahippocampus and OPA; spaces that were found to be more memorable in post-scan interviews showed increased activity in bilateral parahippocampus and the left hippocampus during the movies. For a summary of MRI results showing averaged responses in ROIs across contrasts, see figure 2. Note that while this provides a summary of responses from averaged ROIs, we make inferences using a voxel-wise statistical parametric mapping approach with general linear models (GLM).

**Figure 2:**
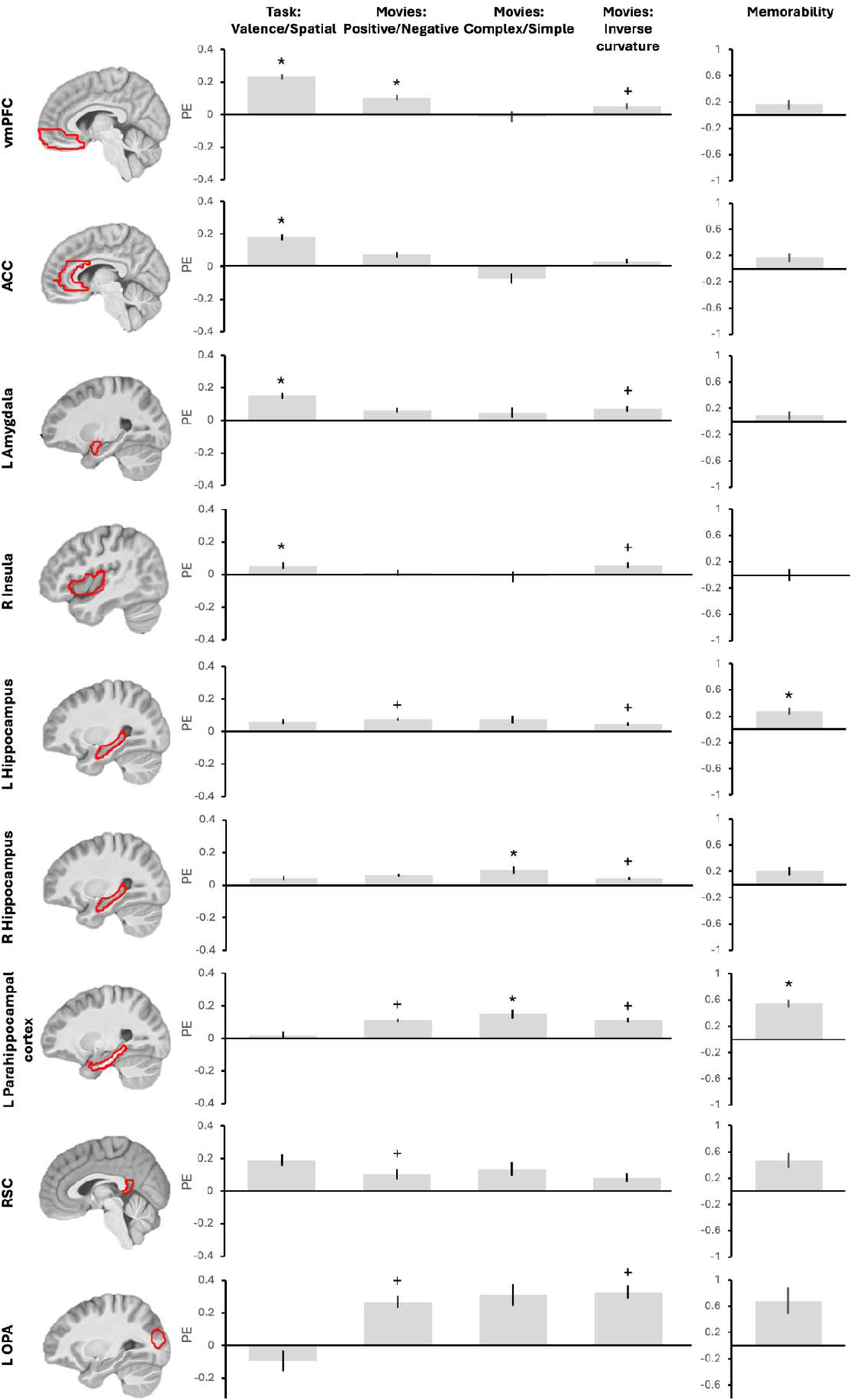
Summary of key MRI results for valence and spatial ROIs in the categorical model. The plots of parameter estimates (PE) provide a summary of the average responses in ROIs for the key contrasts in our analysis (p<.001, minim voxel cluster threshold >5). We present here as a summary an average in regions rather than peak voxel responses, but statistical inference is made at the voxel level using statistical parametric mapping. Valence regions were significantly active in processing positively valenced spaces and valence task focus only. Spatial regions were activated by complex spatial layouts and memorability, but were also activated by positively valenced spaces. Activation survived across valence and spatial regions for inverse curvature (increasing rectilinearity). Brains show ROIs which were anatomically obtained from AAL 3 in the WFU Pick Atlas toolbox, except for the OPA, which was functionally derived by Julian, Fedorenko, Webster, & Kanwisher^75^. Asterisks refer to effects that were predicted and survived small volume correction. Crosses refer to effects that were significant but not initially predicted. Error bars represent standard error of the mean. Hemispheres with significant activation are shown, but similar responses were observed bilaterally in amygdala and insula for task focus, hippocampus for curvature, parahippocampus for complexity and memorability, and both parahippocampus and hippocampus for valence.

All GLMs included the following regressors: button presses, baseline video and novel videos. The primary categorical GLM also included categorical parametric modulators on novel movies for: positive valence, negative valence, simple layout, complex layout, green/blue (GB) space, presence of people, task focus and luminance (latter is non-categorical, mean values). This model was used to carry out categorical contrast analyses e.g. positive vs negative spaces. Non-categorical parametric GLMs were also constructed to incorporate group mean scores for video valence, complexity, fascination, coherence, hominess, unusualness and arousal; computationally derived mean video curvature scores; mean video luminance; and post-scan mean video memorability scores. We report peak level activations from small volume corrections in our ROIs and cluster size. In more exploratory analyses, each contrast was also tested under whole-brain FWE correction (p<.05, minimum voxel cluster threshold >5).

#### Task focus

To examine the effect of task condition, we contrasted valence task focus against spatial layout task focus. Activation for valence task > spatial task survived small volume correction in our valence related ROIs of ACC, vmPFC, bilateral amygdala and bilateral insula (figure 3, table 1). Under FWE correction for whole brain, activation survived in this left rostral ACC (extending into the vmPFC), left precuneus bordering dorsal posterior cingulate cortex, left middle temporal gyrus, bilateral superior temporal gyrus, left superior frontal gyrus/anterior PFC, left angular gyrus, left gyrus rectus (within OFC). We also explored viewing positive vs negative videos specifically under the valence task condition. No valence related ROIs survived small volume correction for negative > positive. For positive > negative, activation survived small volume correction in the vmPFC and left amygdala (table 1). In the same contrast, under FWE correction for the whole brain, activation survived bilaterally in V2, and in both the left parahippocampal gyrus and left fusiform.

**Figure 3.**
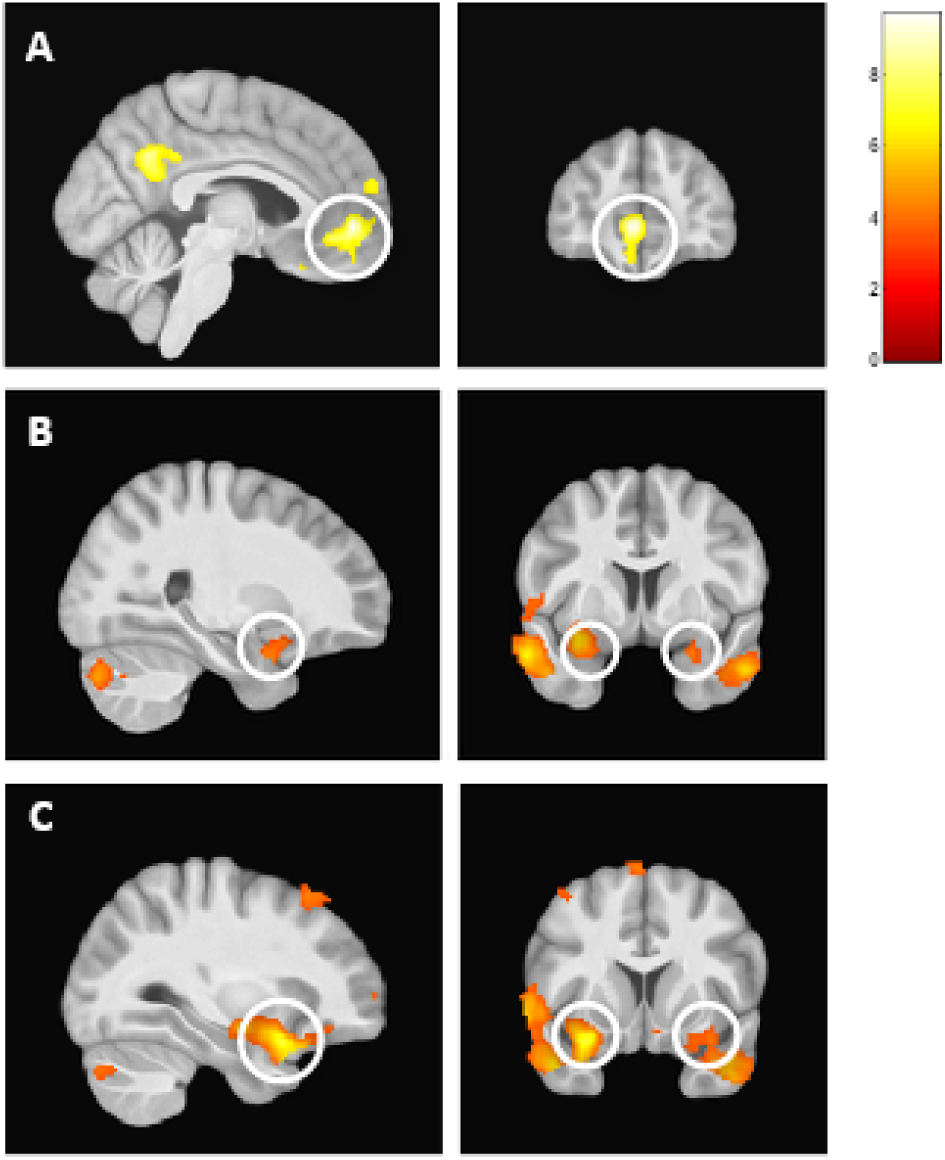
Brain regions more active when viewing movies that require a subsequent valence judgement than a spatial complexity judgement. Regions within the ‘valence network’ were more active during processing of valence task focus > spatial task focus, including: (A) vmPFC (figure thresholded at p=.05 FWE-corrected, minimum 5 voxels), (B) amygdala (thresholded at p=.001 uncorrected, minimum 5 voxels), and (C) insula (thresholded at p=.001 uncorrected, minimum 5 voxels).

**Table 1:** The neuroanatomical correlates of valence related contrasts in affect processing regions. In valence ROIs we report peak activations that survive small volume correction, and cluster size. Coordinates reported in MNI-space.

| Contrast | Area | Peak-level p(FWE) | Z-score | Coordinates | Cluster size |
| --- | --- | --- | --- | --- | --- |
| *Categorical model* |  |  |  |  |  |
| Valence task focus (all videos) > spatial task focus (all videos) | L dorsal ACC | 0.000 | 5.74 | -4, 50, -0 | 1117 |
|  | vmPFC | 0.000 | 6.03 | -4, 52, -4 | 1470 |
|  | R & L amygdala | 0.001 | 4.35 | -28, -8, -12 | 74 |
|  |  | 0.029 | 3.27 | 26, 6, -18 | 4* |
|  | R & L insula | 0.000 | 5.30 | -30, 12, -18 | 194 |
|  |  | 0.007 | 4.23 | 36, 16, -16 | 72 |
| Positive video (valence task focus) > negative video (valence task focus) | vmPFC | 0.014 | 3.99 | 8, 40, -20 | 84 |
|  | L amygdala | 0.002 | 4.04 | -22, -8, -16 | 23 |
| Positive video > negative video | vmPFC | 0.007 | 4.17 | 2, 36, -18 | 230 |
| *Parametric model* |  |  |  |  |  |
| Fascination | R lingual gyrus | 0.000 | 6.55 | 18, -102, -8 | 608 |
|  | vmPFC | 0.056 | 3.57 | 4, 38, -16 | 23 |
| Coherence | L inferior occipital gyrus | 0.000 | 6.41 | -16, -94, -8 | 329 |
| Hominess | L cuneus | 0.001 | 4.53 | -10, -98, 22 | 104 |

\*R Amygdala included below $k > 5$ threshold, shown here to illustrate bilateral responses.

We also contrasted spatial task focus against valence task focus. No activation survived small volume correction in any of our spatial ROIs of hippocampus, OPA, parahippocampal gyrus or RSC, nor under whole-brain FWE correction. We also explored whether our contrasts of complex vs simple altered when looking at these instances specifically under the spatial task condition – as before, no activations survived small volume correction or whole-brain FWE correction. At a more exploratory lower threshold (p<.001, minim voxel cluster threshold >5), for spatial task > valence task we found significant activation in bilateral superior frontal gyrus, corresponding to known localisations of the frontal eye fields (p=.434, Z=4.04, mm=[24, 2, 62], k=445; p=.758, Z=3.75, mm=[−24, −2, 62], k=190)^76,77^.

#### Valence processing

To investigate the processing of valence we first used the categorical GLM and contrasted between negatively valenced videos and positively valenced videos. For negative > positive spaces, no activation survived small volume correction in any of our valence ROIs of ACC, OFC, vmPFC, insula or amygdala, nor with whole-brain FWE correction. We also tested a parametric model that includes group mean scores for valence and complexity. For the contrast negative > positive, no activation survived small volume correction in our valence ROIs of ACC, OFC, vmPFC, insula or amygdala. Nothing survived whole-brain FWE correction.

In the opposing contrast, positive > negative, activation survived small volume correction in the vmPFC in our categorical model (see table 1; figure 4). Activation did not survive small volume correction in the parametric model for this region. With whole-brain FWE correction, the categorical model had significant peak activation in right calcarine, right inferior occipital gyrus, left middle occipital gyrus, bilateral fusiform, but activation was also found bilaterally in the parahippocampal gyrus. Under FWE correction in the parametric model, significant activation was found in the right calcarine and left inferior occipital gyrus.

**Figure 4.**
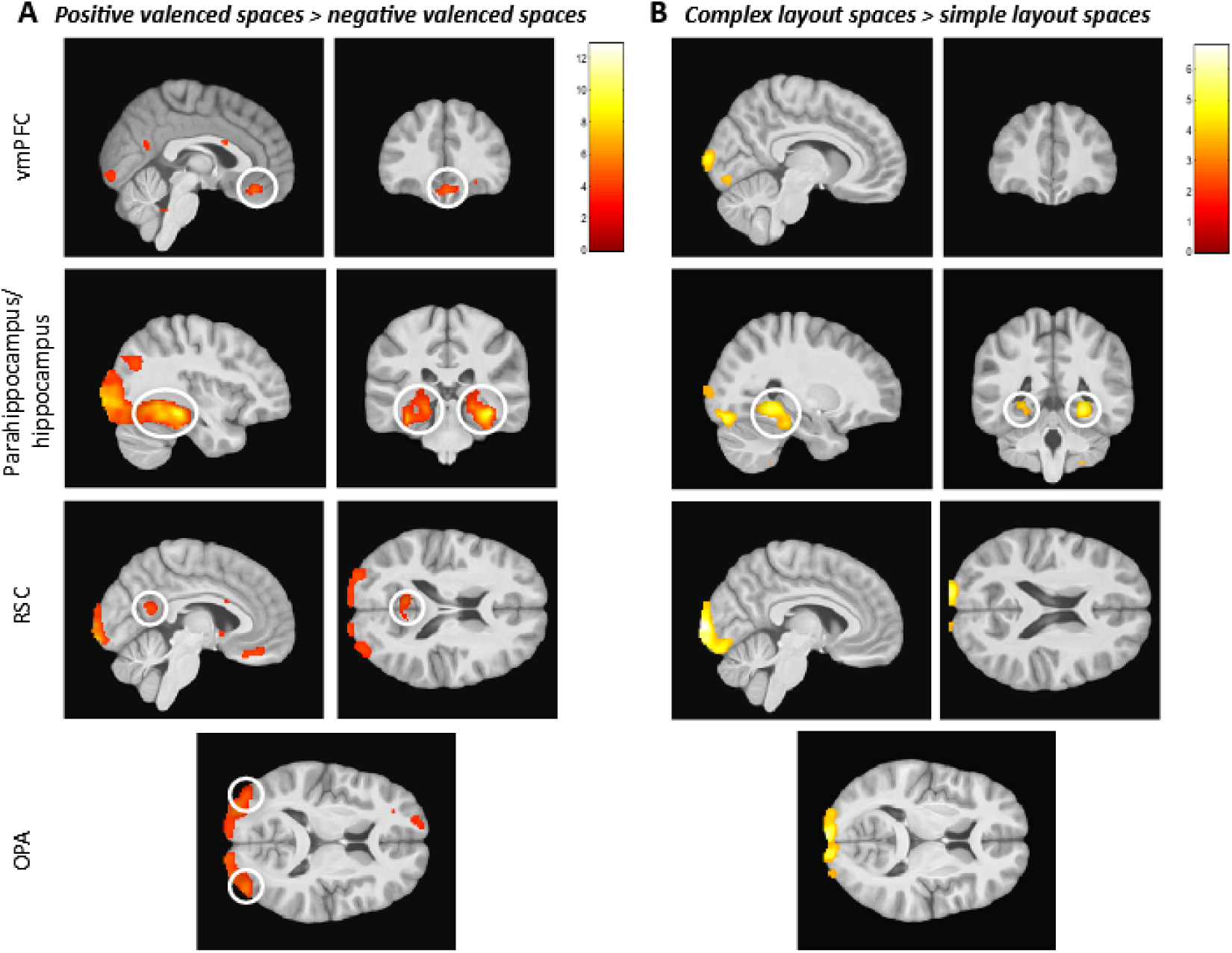
Brain regions more active in spaces that were judged more positive in valence (left) and more complex in their spatial layout (right). (A) During videos that showed positively valenced spaces compared to negatively valenced spaces, there was significant activity in the vmPFC, parahippocampus extending into hippocampus, RSC, and OPA. (B) During videos that showed complex spatial layouts compared to simple spatial layouts, there was significant activity in the parahippocampus extending into the hippocampus, but no significant activity in the vmPFC, RSC, or OPA. Nothing survived in the alternative contrasts of negative > positive or simple > complex. Figures use the categorical model and are thresholded at p=.001 uncorrected, minimum 5 voxels.

To explore other factors associated with aesthetic processing, we re-ran the parametric model replacing group mean valence scores with group mean fascination, coherence or hominess scores (derived from our previous study^4^). Small volume corrections were run across the five valence related ROIs, as well as for the key regions that Coburn et al.^60^ associated with these areas (right lingual gyrus for fascination, left inferior occipital gyrus for coherence, and left cuneus for hominess). For fascination, small volume correction survived in the right lingual gyrus, supporting the findings of Coburn et al.^60^, and vmPFC (table 1). With whole brain FWE correction, activation survived in bilateral calcarine. For coherence, small volume correction survived in the left inferior occipital gyrus, supporting the findings of Coburn et al.^60^ (table 1). Under FWE correction, activation survived in right calcarine and right middle occipital gyrus, as well as in the left inferior occipital gyrus. For hominess, activation survived small volume correction in the left cuneus, once again supporting the hypotheses^60^ (table 1). With whole brain correction, activation survived in the right parahippocampal gyrus, right fusiform, left hippocampus, right calcarine, right cuneus, left superior occipital gyrus, bilateral lingual gyrus.

Given the parahippocampal activation found under FWE correction for positive > negative and hominess, we explored whether valence related activity would survive small volume correction in our spatial ROIs. We observed that in both the categorical and parametric models, under the contrast positive > negative, activation survived small volume correction in our spatial ROIs of hippocampus, OPA and parahippocampal gyrus, and RSC in the categorical model (table 2; figure 4). No activity survived small volume correction in the spatial ROIs for negative > positive. For fascination, coherence and hominess, we observed that activation survived small volume correction in bilateral hippocampus, OPA and bilateral parahippocampal gyrus. Activation also survived in RSC for coherence (table 2). We note that these analyses were exploratory, having identified parahippocampal involvement at FWE correction for whole brain for two contrasts, thus caution should be considered when interpreting these observations.

**Table 2:**
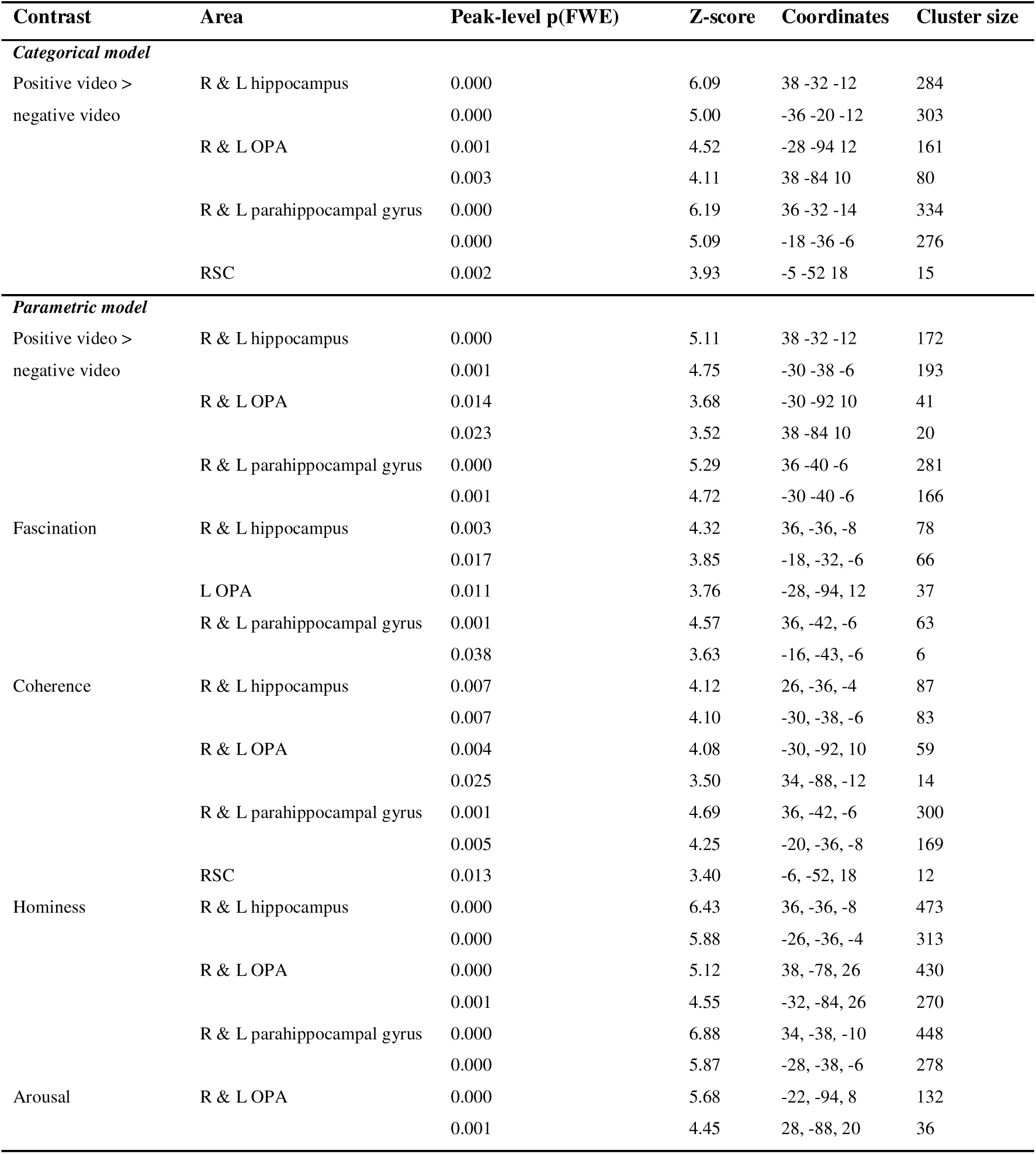
The neuroanatomical correlates of valence related contrasts in spatial processing regions. In spatial ROIs we report peak activations that survive small volume correction, and cluster size. Coordinates reported in MNI-space.

| Contrast | Area | Peak-level p(FWE) | Z-score | Coordinates | Cluster size |
| --- | --- | --- | --- | --- | --- |
| *Categorical model* |  |  |  |  |  |
| Positive video > negative video | R & L hippocampus | 0.000 | 6.09 | 38 -32 -12 | 284 |
|  |  | 0.000 | 5.00 | -36 -20 -12 | 303 |
|  | R & L OPA | 0.001 | 4.52 | -28 -94 12 | 161 |
|  |  | 0.003 | 4.11 | 38 -84 10 | 80 |
|  | R & L parahippocampal gyrus | 0.000 | 6.19 | 36 -32 -14 | 334 |
|  |  | 0.000 | 5.09 | -18 -36 -6 | 276 |
|  | RSC | 0.002 | 3.93 | -5 -52 18 | 15 |
| *Parametric model* |  |  |  |  |  |
| Positive video > negative video | R & L hippocampus | 0.000 | 5.11 | 38 -32 -12 | 172 |
|  |  | 0.001 | 4.75 | -30 -38 -6 | 193 |
|  | R & L OPA | 0.014 | 3.68 | -30 -92 10 | 41 |
|  |  | 0.023 | 3.52 | 38 -84 10 | 20 |
|  | R & L parahippocampal gyrus | 0.000 | 5.29 | 36 -40 -6 | 281 |
|  |  | 0.001 | 4.72 | -30 -40 -6 | 166 |
| Fascination | R & L hippocampus | 0.003 | 4.32 | 36, -36, -8 | 78 |
|  |  | 0.017 | 3.85 | -18, -32, -6 | 66 |
|  | L OPA | 0.011 | 3.76 | -28, -94, 12 | 37 |
|  | R & L parahippocampal gyrus | 0.001 | 4.57 | 36, -42, -6 | 63 |
|  |  | 0.038 | 3.63 | -16, -43, -6 | 6 |
| Coherence | R & L hippocampus | 0.007 | 4.12 | 26, -36, -4 | 87 |
|  |  | 0.007 | 4.10 | -30, -38, -6 | 83 |
|  | R & L OPA | 0.004 | 4.08 | -30, -92, 10 | 59 |
|  |  | 0.025 | 3.50 | 34, -88, -12 | 14 |
|  | R & L parahippocampal gyrus | 0.001 | 4.69 | 36, -42, -6 | 300 |
|  |  | 0.005 | 4.25 | -20, -36, -8 | 169 |
|  | RSC | 0.013 | 3.40 | -6, -52, 18 | 12 |
| Hominess | R & L hippocampus | 0.000 | 6.43 | 36, -36, -8 | 473 |
|  |  | 0.000 | 5.88 | -26, -36, -4 | 313 |
|  | R & L OPA | 0.000 | 5.12 | 38, -78, 26 | 430 |
|  |  | 0.001 | 4.55 | -32, -84, 26 | 270 |
|  | R & L parahippocampal gyrus | 0.000 | 6.88 | 34, -38, -10 | 448 |
|  |  | 0.000 | 5.87 | -28, -38, -6 | 278 |
| Arousal | R & L OPA | 0.000 | 5.68 | -22, -94, 8 | 132 |
|  |  | 0.001 | 4.45 | 28, -88, 20 | 36 |

Though unusualness and arousal ratings were not correlated with valence ratings, their correlation with fascination and complexity ratings^4^ led us to explore these two factors as regressors of interest in a parametric GLM alongside group mean scores for valence, complexity and luminance. We conducted small volume corrections across our valence related ROIs of ACC, OFC, vmPFC, amygdala and insula, as well as our ‘spatial network’ ROIs of hippocampus, OPA, parahippocampal gyrus and RSC. Activations did not survive small volume correction in any of our valence related ROIs for either unusualness or arousal. Activation did however survive in bilateral OPA for arousal (table 2). With whole-brain FWE correction, activation survived for arousal in the right lingual gyrus (the area Coburn et al.^60^ linked to fascination) and bilateral middle temporal gyrus. Nothing survived FWE for whole brain correction for unusualness.

#### Spatial processing

To investigate the processing of spatial complexity, we contrasted videos with simple layouts and videos with complex layouts (simple > complex). In both the categorical and parametric models, no activations survived small volume correction in our spatial ROIs of hippocampus, OPA, parahippocampal gyrus or RSC. Nothing survived whole-brain FWE correction either. In the opposing contrast, complex > simple, activation survived small volume correction in bilateral hippocampus and bilateral parahippocampal gyrus in both models (table 3, figure 4). With whole-brain FWE correction, the categorical model had significant peak activation in the left superior occipital gyrus. Under FWE correction in the parametric model, significant activation was found in the right lingual gyrus bordering the parahippocampus, and the right precuneus/ventral posterior cingulate - part of the RSC.

**Table 3:**
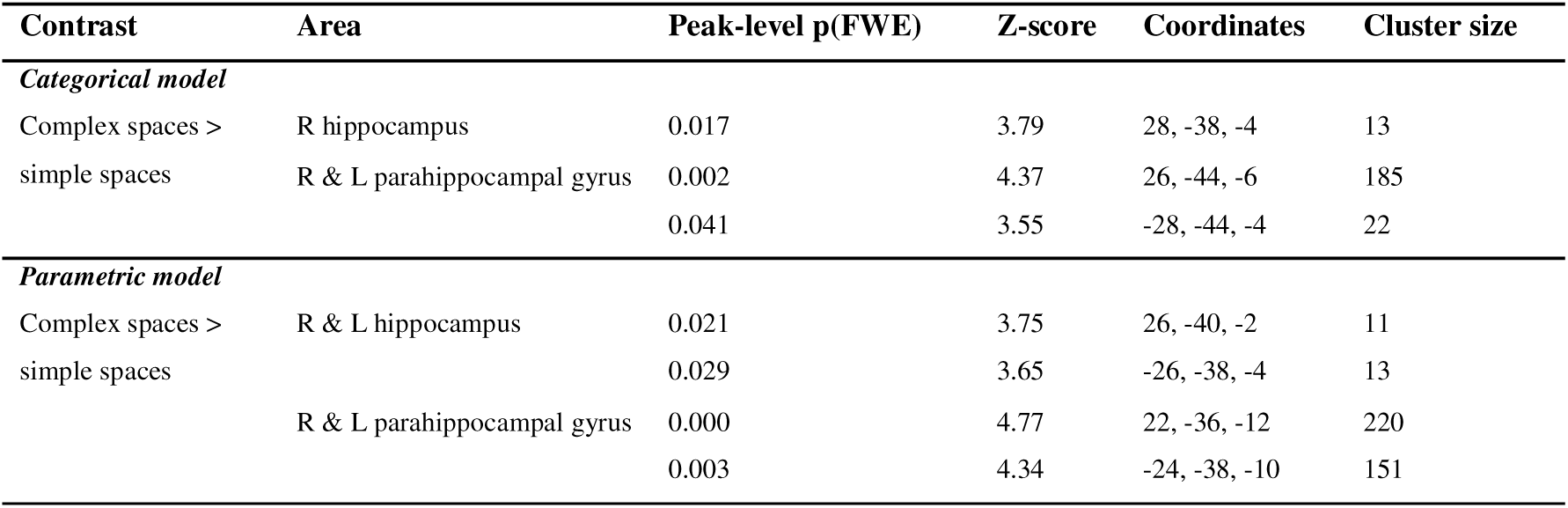
The neuroanatomical correlates of spatial related contrasts in spatial processing regions. In spatial ROIs we report peak activations that survive small volume correction, and cluster size. Coordinates reported in MNI-space.

#### Curvature

With mean values for curvature per video, we could consider which brain regions activated in relation to curvature. See figure 1B for examples of movie stills that show high and low curvature. As described in the methods, mean curvature values were calculated for each video using the MLV Toolbox^78^. Note that curvature was calculated by extracting contours from the scene in each frame, regardless of whether these contours were structural, object-based, textural, etc. We averaged across the curvature scores for each frame, and computed a mean curvature score for each video that reflects all types of contours in the space.

We first explored curvature predicted regions: visual cortex and ACC^39,61^. No activation survived small volume correction in ACC. At an unthresholded correction level (p<.001, k>5), we found activity in left V2 (p=.608, Z=3.89, mm=[−2, −84, −4], k=153) and bilateral associative visual cortex (peaks falling within the OPA). Note that no activation survived FWE for whole brain volume. As curvature can be linked to both spatial geometric features and to aesthetic qualities of space, we performed small volume correction in each of our spatial and valence-related ROIs too. No activation survived small volume correction in any of our ROIs for curvature. We similarly tested for spatial and valence ROIs in the inverse contrast (increasing rectilinearity). Activation survived in three of our valence-related ROIs: amygdala, insula and vmPFC; and across spatial ROIs hippocampus, parahippocampus and OPA (table 4). With FWE correction, activation survived in areas within the right associative visual cortex (middle temporal gyrus) and bilateral middle occipital gyrus.

**Table 4:** The neuroanatomical correlates of curvature in spatial and affect processing regions. In spatial and valence ROIs we report peak activations that survive small volume correction, and cluster size. Coordinates reported in MNI-space.

| Contrast | Area | Peak-level p(FWE) | Z-score | Coordinates | Cluster size |
| --- | --- | --- | --- | --- | --- |
| *Parametric model* |  |  |  |  |  |
| Inverse curvature<br>(increasing<br>rectilinearity) | L amygdala | 0.005 | 3.82 | -30, 2, -22 | 21 |
|  | R Insula | 0.037 | 3.77 | 28, 12, -20 | 13 |
|  | vmPFC | 0.019 | 3.91 | -8, 20, -14 | 17 |
|  | L & R OPA / assoc. visual | 0.000 | 4.98 | 42, -78, 14 | 477 |
|  | cortex | 0.001 | 4.56 | -34, -82, 20 | 269 |
|  | L & R hippocampus | 0.020 | 3.80 | 28, -8, -24 | 82 |
|  |  | 0.025 | 3.72 | -22, -8, -22 | 21 |
|  | L parahippocampus | 0.021 | 3.81 | -30, -28, -18 | 128 |

#### Memorability

We explored whether video memorability activates ROIs of the hippocampus and parahippocampal gyrus - central to memory formation^5,9,79^ - and the amygdala - tied to memory of emotionally arousing information^47,80,81^. To do this, we used two memorability metrics (see methods): free recall scores (from stage 1 of the debrief, in which participants listed the spaces they remembered seeing); and cued recall scores (from stage 2 of the debrief, in which participants were given a visual prompt for each video and then described the rest of the space). Video scores were calculated by counting the number of participants that accurately remembered each video under each memorability task. We explored free and cued video recall scores as parametric modulators tied to movie onsets. For free recall, no activity survived small volume correction in hippocampus, parahippocampal gyrus or amygdala. No activation survived whole-brain FWE correction either. However, activity in the parahippocampus and left hippocampus was higher for movies with higher mean cued recall scores (table 5; figure 5). Under whole-brain FWE correction, cued recall scores activated the left ventral posterior cingulate cortex.

**Figure 5.**
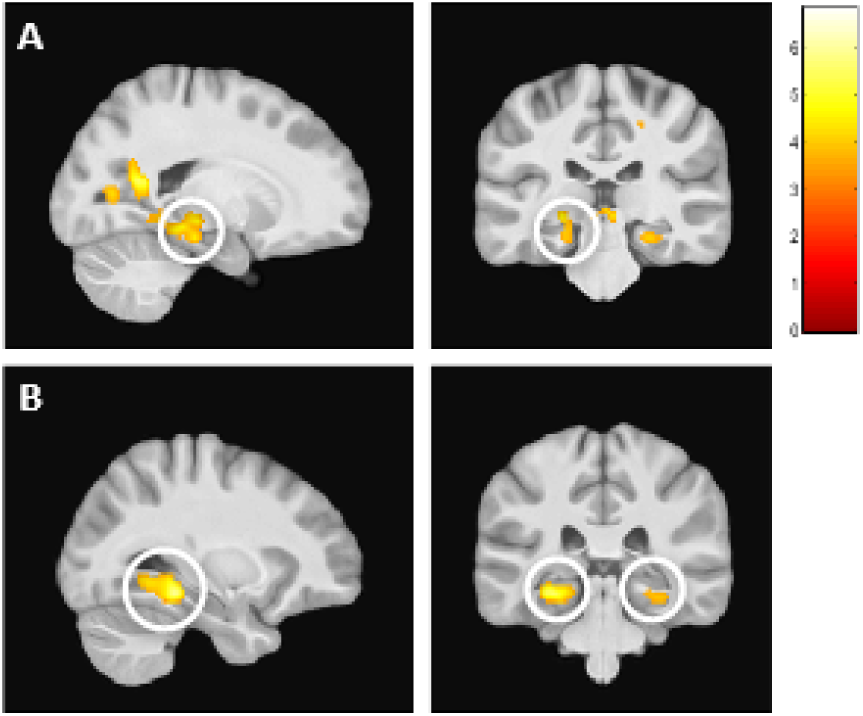
Memorability. Hippocampal-parahippocampal activity increased for videos that are later better remembered during the post-scan debrief. During videos that have high mean cued recall scores (i.e. are recalled correctly post-scan with a 3-second visual reminder), there was significant activity in the (A) left hippocampus and (B) bilateral parahippocampal gyrus. Figures thresholded at p=.001 uncorrected, minimum 5 voxels.

**Table 5:**
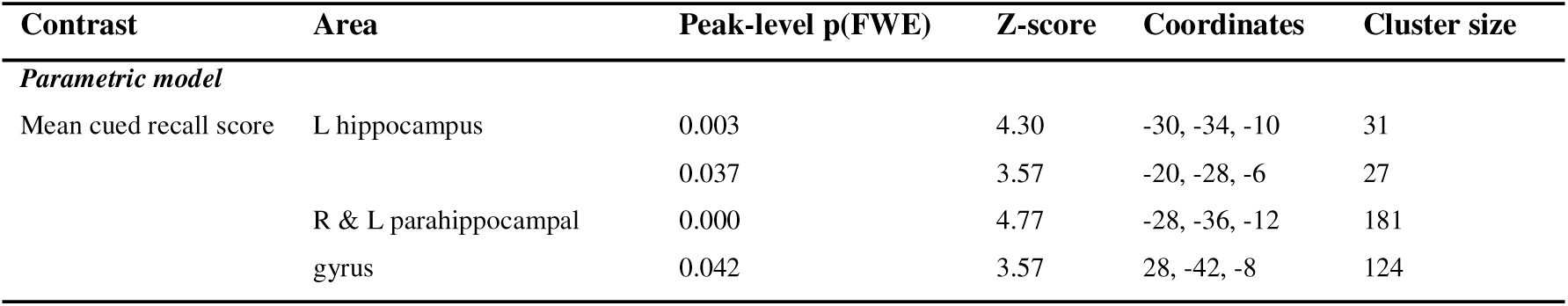
The neuroanatomical correlates of memorability. We report peak activations that survive small volume correction in hippocampal and parahippocampal ROIs and cluster size. Coordinates reported in MNI-space.

### Behavioural results

#### Memorability

To explore memorability, we used two measures (see methods): for each movie we calculated how frequently it was recalled with free recall or with a visual cue (known as cued recall). These scores quantify which spaces were correctly recalled with or without prompt. For correlations between memorability scores and video qualities, see table 6. Video duration did not significantly correlate with any of the memorability metrics (all p>.05).

**Table 6.**
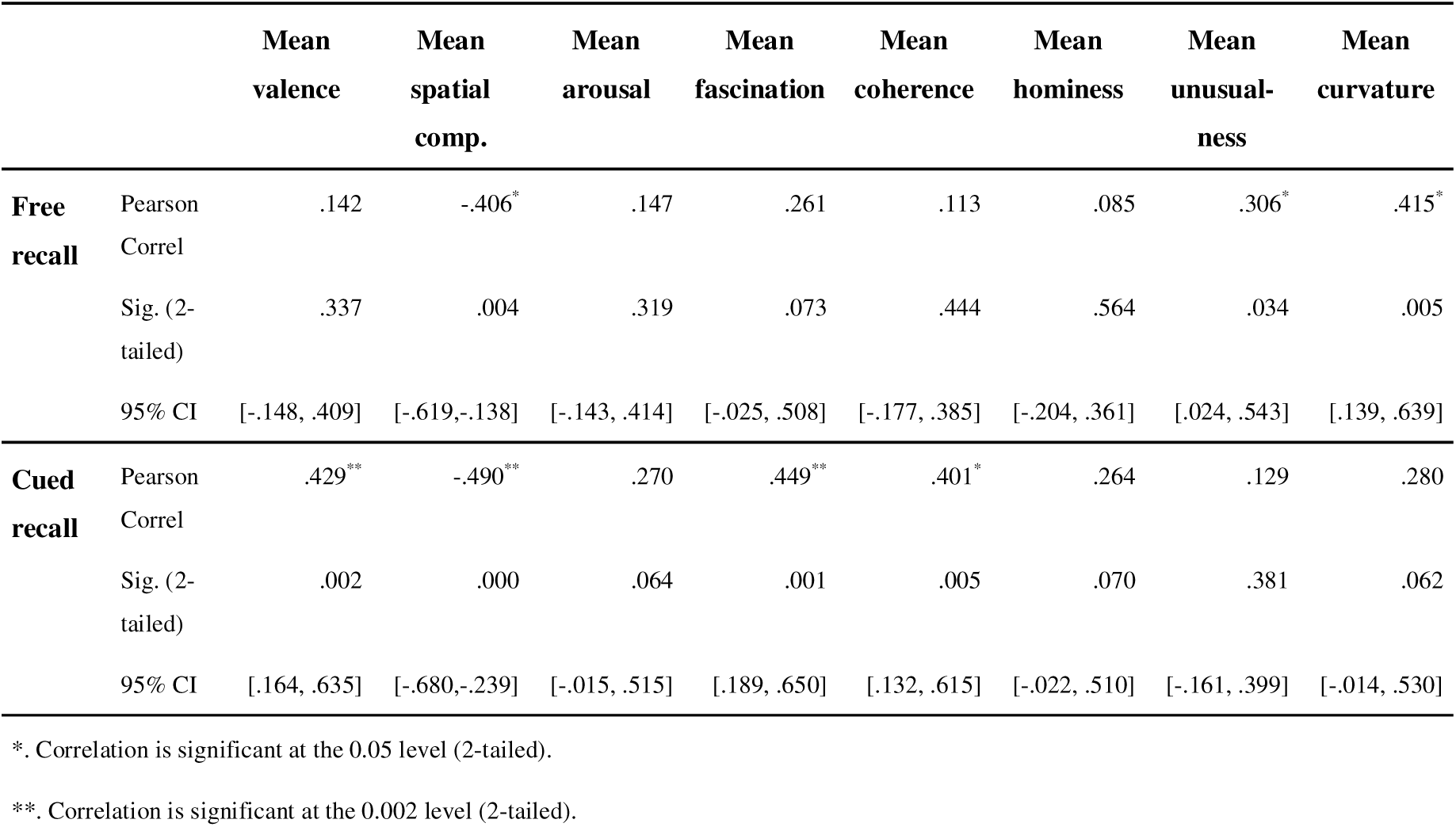
Correlations between memorability scores and video qualities. Correlation analyses were Bonferroni corrected for multiple comparisons, p<.002 (24 comparisons). N.B. mean scores for arousal, fascination, coherence, hominess and unusualness were derived from video analyses in our previous study^4^.

Two-way ANOVAs demonstrated that a video’s valence and complexity did not have a main or interaction effect on free recall scores (p>.05), but that complexity had a main effect on correct cued recall scores (F(1, 44)=4.449, p=.041). Simple spaces were more likely to be remembered correctly in cued recall than complex spaces. We also used two-way repeated measure ANOVAs to explore whether free and cued recall scores differed within participants, according to valence and complexity. To do this, for each participant we calculated the frequency that e.g. a positive simple video was recalled correctly and divided that by the total number of positive simple videos in the set. This gave subject-specific scores for the memorability of positive simple, positive complex, negative simple and negative complex spaces. There was a statistically significant interaction between the effects of valence and complexity on a participant’s free recall (F(1,31)=5.727, p=.023) and cued recall (F(1,31)=28.914, p=.000). Paired t-tests indicated that for both free and cued recall, positive simple spaces had higher memorability than positive complex, negative simple or negative complex spaces (p<.05). Finally, we explored whether task focus impacted memorability by comparing recall performance across the two participant groups that experienced different tasks for each movie (e.g. PG1: valence, spatial, valence… vs PG2: spatial, valence, spatial…). Free recall scores for both groups correlated strongly and positively (r=.648), as did cued recall scores (r=.706). Two sample t-tests showed no significant difference between free recall scores or between cued recall scores (p>.05). This indicates that task focus did not impact memorability of videos.

#### Valence and complexity

Participants in the fMRI scanner were asked to rate either the valence or the spatial complexity of each space while watching a series of first-person videos walking through architectural spaces (see figure 1). Mean valence and complexity scores for each video were calculated using the MRI group’s in-scanner ratings and videos were subsequently categorised as positive or negative, and simple or complex. We could then compare these mean video scores with the mean scores derived on the same videos in our previous survey^4^. All MRI ratings were converted into a 1-9 scale for comparison. Overall, 84% of our binary video categorisations (positive or negative, simple or complex) remained the same across both studies. One video switched in valence (positive>negative) and 14 switched in complexity (11 complex>simple, and 3 simple>complex).

There was a strong positive correlation of mean valence scores (r=.971) between the current study and our prior survey, and a moderately strong, positive correlation between the current study and the prior survey’s mean complexity scores for each video (r=.644). This suggests that participants in both studies overarchingly evaluated our videos similarly. Independent sample two-tailed t-tests (equal variances not assumed) confirmed that mean valence scores were not significantly different between the two study groups (p>.05), but that mean complexity scores were significantly different between the two study groups [*t* (72.485)=−2.156, *p*=.034]. The variance in mean video complexity scores was smaller in the survey group (s^2^ = .872) than in the MRI group (s^2^ = 2.95); complexity ratings were less spread in our original study.

In order to check whether these correlations were driven by the performance of a particular participant group (PG1 or PG2), we also examined each group separately. We found a strong positive correlation between the mean video scores derived from PG1 and their equivalent scores from the prior survey (r=.856), and a strong positive correlation between the mean video scores derived by PG2 and their equivalent scores from the prior survey (r=.907). Independent sample two-tailed t-tests (equal variances not assumed) also confirmed that mean video scores did not differ between the survey and PG1, nor the survey and PG2 (all p>.05), suggesting that MRI participants performed largely in-line with participants from the survey. For the relationship between our mean valence and complexity scores and some key qualities of interest (e.g. fascination, calculated from Gregorians et al.^4^), see table 7.

**Table 7.** Correlations between mean valence and complexity scores and key qualities of interest. We compared correlations for mean valence and complexity scores derived from this MRI study and from our previous survey study. Correlation analyses were Bonferroni corrected for multiple comparisons, p<.002 (29 comparisons). CI = confidence interval.

|  |  | Mean<br>arousal | Mean<br>fascination | Mean<br>coherence | Mean<br>hominess | Mean<br>unusualness | Mean<br>curvature |
| --- | --- | --- | --- | --- | --- | --- | --- |
| **Mean valence**<br>(fMRI study) | Pearson Correl | .110 | .839** | .893** | .809** | .094 | -.117 |
|  | Sig. (2-tailed) | .456 | .000 | .000 | .000 | .524 | .442 |
|  | 95% CI | [-.180,.382] | [.728,.907] | [.816,.939] | [.682,.889] | [-.195,.369] | [-.397,.182] |
| **Mean valence**<br>(prior survey) | Pearson Correl | .202 | .840** | .887** | .806** | .108 | -.149 |
|  | Sig. (2-tailed) | .088 | .000 | .000 | .000 | .368 | .328 |
|  | 95% CI | [-.0311,.414] | [.755,.897] | [.825,.928] | [.706,.874] | [-.127,.331] | [-.424,.151] |
| **Mean spatial**<br>**complexity**<br>(fMRI study) | Pearson Correl | .003 | -.190 | -.176 | -.180 | -.194 | -.312* |
|  | Sig. (2-tailed) | .983 | .196 | .231 | .222 | .186 | .037 |
|  | 95% CI | [-.281,.287] | [-.450,.099] | [-.438,.114] | [-.441,.110] | [-.453,.095] | [-.555,.020] |
| **Mean spatial**<br>**complexity**<br>(prior survey) | Pearson Correl | .405** | .478** | .160 | .099 | .335* | -.168 |
|  | Sig. (2-tailed) | .000 | .000 | .181 | .409 | .004 | .271 |
|  | 95% CI | [.191,.582] | [.277,.639] | [-.744,.378] | [-.136,.323] | [.112,.526] | [-.440,.132] |
| **Mean**<br>**curvature** | Pearson Correl | .170 | .055 | -.250 | -.099 | .365* | 1 |
|  | Sig. (2-tailed) | .266 | .719 | .098 | .519 | .014 |  |
|  | 95% CI | [-.130,.441] | [-.242,.343] | [-.506,.047] | [-.381,.201] | [.080,.595] |  |

\* Correlation is significant at the 0.05 level (2-tailed), but does not meet threshold for significance.
\*\* Correlation is significant at the 0.002 level (2-tailed).

## Discussion

Using first-person view videos of journeys through buildings and cities, we examined how brain regions involved in spatial mapping – hippocampus, OPA, parahippocampus and RSC – and brain regions involved in affective processing – ACC, OFC, vmPFC, insula and amygdala – may be activated during architectural experience. We found that valence-related brain regions were activated when participants were tasked to track affective, as opposed to spatial, aspects of the environment. Consistent with prior predictions we found that experiencing complex spaces activated the hippocampus and the parahippocampus, and that experiencing positively-valenced spaces activated the vmPFC. Although previous studies have suggested a distinct activation of anterior temporal and posterior medial processing networks^70^, we found that journeys through environments with positive valence activated not only areas in our valence-related regions but also activated spatial processing regions - hippocampus, OPA, parahippocampus and RSC.

When participants were asked to evaluate how pleasant a space was, all of our key valence-related brain regions were found to be more active. And, as anticipated, we found that positively valenced spaces activated the vmPFC. This finding is consistent with the view that the vmPFC plays a role in value judgments and signalling positive valence across different stimuli^40–43^. Our finding that the response was categorical, responding to positive > negative movies, rather than tracking the valence of each movie, may suggest that, at least for these stimuli, it was engaged in a more binary evaluation process (is this place pleasant or unpleasant?) rather than tracking the level of valence in the stimuli. This is consistent with prior studies in which participants judged valence as categorically positive or negative^82,83^. Notably our stimuli were designed to exist in two categories: either positive or negative. Thus, there may have been insufficient variation in valence to facilitate parametric tracking of valence in our study. Whilst we only found vmPFC activation in relation to positive spaces, prior studies involving valence stimuli have had mixed results, with vmPFC activation reflecting positive, negative, or both ends of the valence spectrum^82,83^. It is possible that for architectural spaces the positive valence provides greater overall engagement with the space, attraction to features, and greater demand on consideration of the space compared with the negative spaces. Future research will be useful to explore more around the phenomenological aspects of positive compared to negative spaces.

Beyond the vmPFC, we were surprised to find that activity in the parahippocampal region was significantly more active in positive vs negative spaces, when correcting for whole brain volume. This prompted us to explore whether positive spaces would drive increased activity across the regions associated with spatial processing, which indeed was the case. Results indicated that positive spaces activated all our spatial-related ROIs. Following prior work^70^, we had not anticipated overlap between networks involved in aesthetic and spatial mapping, but our results indicate that positive valence spaces engage both groups of brain regions. As previously mentioned, whilst these spatial regions are traditionally associated with scene processing and spatial memory, there is some prior evidence that points to their role in valence processing^52–59,84^. Our finding of increased RSC activity for positive spaces is broadly consistent with evidence that this region is more active when viewing neutral scenes rather than threatening scenes^84^. These findings perhaps support the notion that high affective value may provide key contextual cues that help situate objects within spaces, facilitating emotion processing and generating expectations about the environment^85^. Whilst these regions were not predicted for valence processing, they perhaps could have been given the prior evidence. However, further work is needed to deduce what drives this activity in spatial processing regions; whether these responses are borne from visual properties of the stimuli themselves; and whether widespread activity in visual areas covers hippocampal and parahippocampal activity. Whilst efforts were made to control for visual properties of the stimuli (including luminance, optic flow, hue and saturation - see methods), the selection of videos could be further controlled with more fine grain parameters, to therefore systematically separate visual properties of the stimuli from aesthetic qualities. However, this is difficult to entirely control for with ecological stimuli. Nonetheless, these findings add insight to the question of how ‘spatial regions’ relate to valence processing, particularly in the architectural domain; whilst previous studies have considered emotionally salient experiences in space, our work considers the experience of the space itself.

Another key finding is that complex spaces activated parts of our spatial network in all of our models. This supports the hypothesis that processing of more navigational choice points, i.e. higher complexity, would increase the load on path planning and scene-processing regions^86^. Whilst some studies have found hippocampal and RSC activation to increase in simple rather than complex spaces - in that simpler spaces may be easier to process for navigation - this concerns active navigation, where decisions have to be made^26^. Our study involved passive viewing, where the participant made no navigational decisions; the PPA and RSC are known to respond strongly during passive viewing^20,87^.

Previous work has found that fascination, coherence and hominess strongly and positively relate to valence^4,60^. Here, we confirm this result and expand on this line of work. Firstly, we are able to replicate the neural underpinnings of these three aesthetic dimensions found by Coburn and colleagues^60^, but we do so using dynamic rather than static stimuli. Secondly, we were able to show that this association holds irrespective of task focus. As with valence, all three dimensions also activated spatial regions including the hippocampus, OPA, parahippocampus, and, exclusively for coherence, the RSC. Notably, while we were able to test whether different properties did engage regions predicted to support spatial and aesthetic processing, our movies were not designed to distinguish responses between valence, fascination, coherence and hominess, as they were correlated. Curvature of spaces is an interesting metric that relates to both geometric processing and valence, in that people tend to prefer curved objects and spaces^67,68,88^. We calculated mean curvature scores for each video, and found that curvature correlated with activity in the visual cortex, consistent with prior findings, but at the uncorrected threshold^39,61^. For inverse curvature (rectilinear spaces) we found activity increased in both our valence and spatial processing brain regions. Notably, this included activation in the amygdala, which had been previously hypothesised to correlate with rectilinearity, but had not been empirically supported in the architectural context^39^. Amygdala activation has been empirically tied to rectilinearity in other domains, such as for ratings on sharp versus curved-contour everyday objects^67^. Our behavioural results showed valence was not directly correlated with curvature. Rather the more curvature in a space, the simpler and the more unusual it tended to be rated. A recent study by Vartanian et al.^61^ similarly found that curvature scores calculated from images of spaces (as we did here) did not correlate with aesthetic pleasure, whereas perceived curvature correlated with pleasure. We explored the effect of curvature as it is a well-studied architectural quality with empirical hypotheses to test, but there is scope to extend this type of enquiry further. Videos can be analysed to explore changes in other design features such as openness, lines of sight, ceiling height or enclosure, which have all been tied to aesthetic or spatial processing and behaviour^60,62,89^. Future research exploring curvature along with other properties of space will be useful to help understand the impacts of design properties.

Regarding memorability, we found that changing complexity may be a key driver in our study. Simpler spaces were more likely to be remembered correctly in free and cued recall. This supports prior studies showing that more complex spaces lead to poorer wayfinding, mental representation and memory for places^90,91^. In line with findings that emotionally charged information is more memorable^81,92,93^, we found that valence held a moderate positive correlation with cued recall scores. The literature on whether positive or negative valence increases memory differs. Some studies find that negative valence enhances memory, including sensory contextual detail^94–97^, whilst others find that positive valence improves memory, and that negative valence has no impact over memory for spatial and temporal contexts^98,99^. Our finding that positive spaces correlated with successful cued recall may be explained by positing that encoding of these positive scenes taps into self-referential processing or autobiographical memory, with participants connecting positive scenes to past experiences, which improves their memorability^99^. In addition to simple layouts, we found that unusualness and curvature also correlated with increased free recall scores, and that pleasantness, fascination and coherence also correlated with increased cued recall scoring. All of these factors had been linked to memorability previously in the literature, often with contradictory results^100–104^. With regards to brain regions, we found that cued recall scores activated key memory encoding regions of the parahippocampus and hippocampus, supporting the hypothesis that memorability is related to successful encoding in the hippocampal-parahippocampal network^105^.

One limitation of our approach is that, while incorporating a dynamic component, we still lack the multimodality that is characteristic of real-world architectural experiences. Similarly, we have participants rating each space only at the end of each video. Whilst the use of singular post-stimuli ratings has been validated^106,107^, it would be interesting to track the dynamic changes in pleasantness or complexity within a video. Timestamped increases and decreases in pleasantness or complexity can be modelled in our existing GLMs, to give a more fine-grained analysis of changing dynamics. By introducing the measure of curvature to this study, we began to explore the relationship between aesthetic coding, spatial coding, and design properties of the environments. As mentioned, we anticipate that this will be a fruitful line of enquiry that has a multitude of opportunities, exploring other design properties. Future research may also consider individual differences including personality types, architectural training, and rural vs city upbringings, as all have been found to impact navigational performance, spatial processing, or affective response^108–111^. As for our finding that positive spaces activated the spatial network, a potential explanation is that these spaces are more memorable. Further analysis is needed to understand this root cause: whether pleasant spaces can propel scene processing. This could be tested by considering how, for example, the valence of a space impacts wayfinding performance. Finally, an exploratory analysis found activation in bilateral frontal eye fields for the spatial complexity task focus, which suggests that participants might be looking more extensively and rapidly around the screen to assess the complexity of the space. Future analysis of eye tracking data will be useful to test this prediction^112^.

Our study explored the neural correlates associated with evaluating the valence and spatial layout complexity of architectural spaces, using a dataset of first-person videos of trajectories through different built environments. As anticipated, we found that key valence-related regions activated in relation to positive spaces compared to negative spaces and that key spatial regions activated in relation to complex rather than simple spaces. Most notably, we found these spatial mapping regions to also activate for positive compared to negative spaces, suggesting that positive valence processing engages networks supporting both aesthetic/affective and spatial mapping. These findings open further lines of enquiry, but also shed light on what factors of the built environment might impact architectural experience, how they relate to one another, and how different brain regions process them. Though spatial and aesthetic mapping of environments have largely remained in separate fields of enquiry, our work suggests that they may in fact be highly entwined.

## Methods

### Recruitment

Participants were recruited via UCL SONA and by word of mouth. Interested participants filled out a pre-screening survey to ensure that they did not meet the exclusion criteria, including if they: were architects or architecture students; prone to car-sickness or nausea; claustrophobic; would be uncomfortable lying on their back for ∼1hr; left-handed; or if they would not pass the MRI safety checks. Participants who successfully passed pre-screening were invited to participate. Ethics approval was given by the first author’s institution prior to this initial recruitment. As is discussed later, this resulted in complete scanning of 34 participants, of which 26 were included after quality checks using MRIQC (see Analysis section). Recruitment targets were derived from previous fMRI studies on architectural experience, which range from 18-23 participants^39,62,113^.

### Pre-study training

In the week prior to their MRI scan, participants were sent an online questionnaire which included an information sheet detailing all stages of the experiment, consent form and demographic questions. Participants also completed the Navigational Strategy Questionnaire (NSQ^73^), and Spatial Anxiety Questionnaire (SAQ^74^). Participants were placed in one of two groups according to demographics (PG1 or PG2). Independent sample t-tests confirm that average NSQ and SAQ scores are not significantly different between the two participant groups (p>.05). We do not analyse these data further in this article.

The online form then took participants through a training exercise to familiarise themselves with the task they would complete in the scanner, and the terms and symbols they would need to understand. Icons for valence and complexity were designed so that participants could quickly and effectively identify the task condition during the experiment. Both terms were also defined; valence referring to how pleasant or unpleasant a space is (on a 1-5 scale, where 1 indicates very unpleasant and 5 very pleasant), and spatial layout complexity referring to how simple or complex they found the space (on a 1-5 scale, where 1 indicates not at all complex, and 5 extremely complex). As the less colloquial term, additional information was given to define spatial complexity as the route options and spaces you might be able to walk through from your current position – i.e. the spaces that are directly available to you, not to be confused with the imagined layout of the rest of the building, or visual complexity. Reference images were finally used to visually describe these terms (though it was clarified that these images did not necessarily denote the extreme ends of the scales).

Participants then went through two practice runs of the task with two videos that are not featured in the main experiment. After giving their ratings for these videos, participants were told the average scores that these videos had received in our previous behavioural study^4^, and were asked to confirm if this made sense to them; this helped identify if any participants needed further explanation on the task structure or definitions.

Finally, participants were introduced to a ‘baseline video’. They were informed that they would see this video multiple times in the main experiment, and that we wanted them to feel comfortable and familiar with the space. Though they must watch the video, they would not need to assess it for pleasantness or complexity – they are asked to always rate this space as neutral (3 on the 1-5 scale).

### Stimuli selection

We previously curated a database of 63 videos that varied in valence and complexity^4^. In order to keep overall scanning time under 55 minutes, 48 videos were included in this study. The 48 videos were chosen and distributed into four blocks of 12 videos. Using data gathered on these videos in our previous study^4^, blocks were evenly distributed (to the greatest extent possible) in valence, arousal, complexity, building types, presence of people, and presence of GB space. Videos with GB space were further classified as having either pleasant or unpleasant GB spaces by the researcher (six videos could not be classified as having distinctly pleasant or unpleasant GB spaces).

### MRI acquisition parameters

Scanning was conducted at the Birkbeck-UCL Centre for Neuroimaging (BUCNI) using a 3-T Siemens Prisma MRI scanner with a 32-channel head coil. Each scanning session consisted of four runs, totalling approximately 55 minutes. A multi-band sequence was used (repetition time [TR] = 1450msec, echo time [TE] = 35 msec, flip angle = 70°). The scans were whole-brain (72 slices) with a multi-band acceleration of 4, slice thickness of 2 mm, spacing between slices of 2 mm, resolution / voxel size of 2×2×2 mm, field of view of 212 mm, and phase encoding A >> P. A standard T1-weighted high-resolution structural scan (MPRAGE) was acquired before the first run (TR = 2300 msec, TE = 30 msec, 1×1×1 mm resolution), and a fieldmap after the second run. A top-up sequence was also performed at the end of the scanning session with reverse-phase encoding. Ear plugs were used to reduce noise, and foam padding on each side of the head to minimise head movements. Participants viewed the task stimuli via a mirrored rear projector screen and were given a right-hand button box in order to give their ratings during the task. Heart rate, breath rate and eye tracking were also collected throughout the experiment, though this data goes beyond the scope of this paper.

### Task structure

On arrival at the scanning unit, participants were once again given an information sheet and consent form, and were taken through the MRI scanning safety checks and task structure. They were shown the baseline video once more, and completed the Positive and Negative Affect Schedule (PANAS^72^) to gather an indication of their current emotive state (not analysed in the current study).

The overall in-scanner task structure is detailed in figure 1. At the start of each run the participant saw a fixation cross for 4.35s, followed by a 12s warm-up video that walked them down a straight corridor (N.B. this is not the baseline video). This video was used to focus the participant before the experiment began. Then, the task cycle commenced. A screen was shown indicating whether the participant would need to focus on the valence or spatial complexity of the following video (2-4s, fixed randomisation across participants). The video then played with the task icon displayed at the top of the screen as a reminder of whether they should be focusing on valence or complexity (31-40s).

The participant was then asked to rate the video for the given task (valence or complexity) on a 1-5 scale using their right-hand button box. First responses were recorded, and the screen moved on after 7.25s. This task cycle then repeated for 6 more videos, before reaching the halfway break point. An image of a waiting room was displayed for 10s, after which the task resumed with the fixation cross and the same warm-up video as before. The above process repeated with 7 new videos. At the end of the block, the participant got another break, and was asked whether they were happy to continue. This marked the end of a block and run. Total block length was ∼12 minutes. The task structure was piloted and iterated a series of times outside of the scanner before in-scanner piloting and subsequent data collection commenced.

Within each block, videos alternated in positive or negative valence, and task focus alternated between valence or spatial layout complexity. Block order was fixed as was video order, however task focus was inverted between the two participant groups for partial counterbalancing. Therefore, if group PG1 were under the valence task condition for the first video in block A, group PG2 would experience that same video under the spatial complexity task condition.

### Post scan

After scanning was complete, participants had a 10-minute break, and then had a debrief interview consisting of three stages. The debrief session was audio recorded, and notes made by the researcher.

**Stage 1: free recall test.** Firstly, they were asked to list as many spaces as they could remember. This section lasted a minimum of 5 minutes but could extend further.

**Stage 2: cued recall test.** In the second stage, participants watched back the first 3s of each video. They were asked to succinctly talk through the rest of the route – where they went, what they saw, or how they felt. They could also express if a space looked completely unfamiliar to them, or if it looked familiar but they could not recall any details.

**Stage 3: final questions.** Finally, participants were asked: whether they recognised or were familiar with any of the spaces; how engaged or immersed they were when watching the videos in the scanner (qualitative response and 1-10 rating); whether they felt particularly nauseous at any point; and whether they felt their reaction to the baseline videos altered over time.

### Analysis

#### Video characteristic analysis

Mean values for luminance, optic flow, hue and saturation were calculated for each video to understand whether low level visual features needed to be accounted for in further analyses. One-way ANOVAs indicated that mean luminance was statistically different between positive simple, positive complex, negative simple and negative complex videos (F(3,44)=4.201, p=.011). Tukey’s HSD test for multiple comparisons found that luminance was higher in positive simple videos than negative simple (p=.044, 95% C.I.=[.3556, 37.4977]) or negative complex videos (p=.017, 95% C.I.=[3.0147, 41.0646]). Mean optic flow, hue and saturation were not statistically different between the four video groups (p>.05).

Chi-square tests for independence also confirmed that there was no significant association between mean valence and presence of people or GB space, nor mean complexity and presence of people or GB space (p>.05). Correlations between luminance and people (r= −.005) and luminance and GB space (r=.142) were also not significant and weak.

With these results, we could deduce that luminance was the only video feature that would need to be considered in further analyses centring on valence and complexity.

Finally, mean curvature values were calculated for each video using the MLV Toolbox^78^ (N.B. curvature could not be calculated for 3 videos). Curvature is defined as ‘the change in angle per unit length’, and calculated according to how line segments change in orientation in proportion to the segment length. Therefore higher values indicate a higher rate of change in orientation along a contour. It is important to note that curvature was calculated by extracting contours from the scene, regardless of whether these contours were structural, object-based, textural, etc. Therefore the metric of curvature reflects an average across all types of contours in the space. A one-way ANOVA indicated that mean curvature was not statistically different between positive simple, positive complex, negative simple and negative complex videos (p>.05).

#### Behavioural analysis: valence and complexity ratings

34 participants completed the study and rated each video for either its valence or complexity while in the MRI scanner. Each participant’s 48 video ratings were first considered in relation to the mean rating that those same videos had received in our previous study^4^, in order to evaluate how our participant pools compared. If a participant rated >66% videos in line with the prior survey results (i.e. rated a video as positive in valence, which was also deemed positively valenced in the prior survey), they were considered to have responded largely in-line with the participant pool that generated the video metrics in our previous work. Four participants did not meet this criterion, and so had further data checks. We first checked whether their ratings for the most and least valenced/complex places matched the survey results. Only one of these four participants failed to meet these criteria, and their data was not included in analysis, as we considered them an exceptional outlier.

With the 33 included participants’ data (Group 1: M=8, F=8; 18-24yrs=9, 25-34yrs=7; Group 2: M=10, F=7; 18-24yrs=16, 25-34yrs=1), mean valence and complexity ratings were calculated for each video. Analyses in this paper were carried out at the video level using these mean scores; correlations and t-tests were used to compare these new mean video ratings with those of the previous survey (e.g. fascination, coherence and hominess).

#### Memorability analysis

Memorability had two components: free recall (stage 1 of the debrief), and cued recall (stage 2 of the debrief). Free recall scores were calculated at the individual and video level by A) calculating the percentage of positive simple, positive complex, negative simple and negative complex spaces each participant remembered in stage 1 of the debrief, and B) counting the number of participants that remembered each video in stage 1. A space was deemed to be ‘remembered’ if the participant could describe the space with enough identifiable detail, which could not apply to any of the other spaces (e.g. the palace with red walls and gold chairs > the palace). On occasion, it was unclear which space a participant was describing, or the researcher identified that they were merging memories of different spaces – these descriptions were discounted, and on average this totalled 1 video per participant.

The cued recall data from stage 2 of the post-scan debrief was first coded: 0 = video not recalled; 1 = video somewhat recalled but without detail; 2 = video seemingly recalled, but participant is describing the wrong space; 3 = video is correctly recalled. See example statements and ratings in table 8.

**Table 8.**
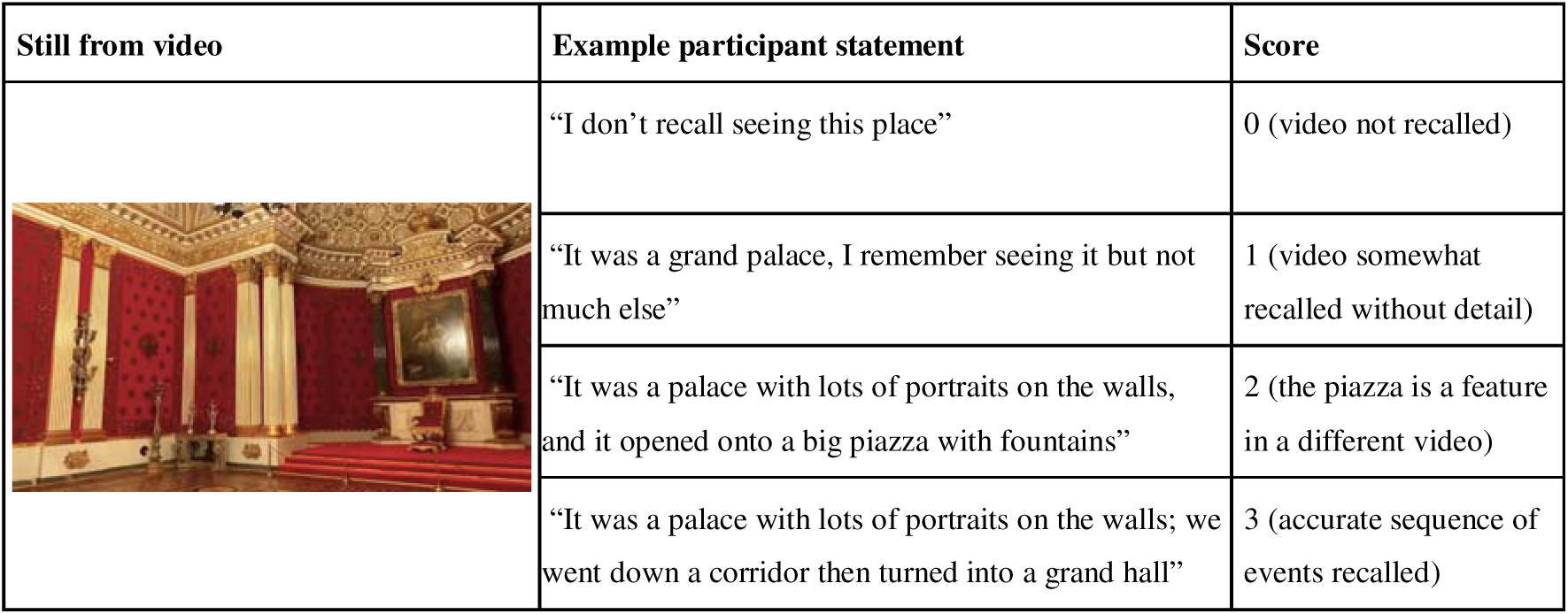
Example illustrative statements for coding the cued recall memorability task in stage 2 of the debrief.

| Still from video | Example participant statement | Score |
| --- | --- | --- |
|  | “I don’t recall seeing this place” | 0 (video not recalled) |
|  | “It was a grand palace, I remember seeing it but not much else” | 1 (video somewhat recalled without detail) |
|  | “It was a palace with lots of portraits on the walls, and it opened onto a big piazza with fountains” | 2 (the piazza is a feature in a different video) |
|  | “It was a palace with lots of portraits on the walls; we went down a corridor then turned into a grand hall” | 3 (accurate sequence of events recalled) |

Cued recall scores were then calculated at the participant and video level by A) calculating the percentage of positive simple, positive complex, negative simple and negative complex spaces that each participant did not recall, somewhat recalled, wrongly recalled, and correctly recalled in stage 2 of the post-scan debrief, and B) counting the number of participants that did not recall, somewhat recalled, wrongly recalled, and correctly recalled each video in stage 2. Therefore, cued recall consisted of four scores at both the participant and video level.

Stage 3 of the post-scan debrief was used to ensure our data was robust. Participants overwhelmingly did not recognise any of the spaces, felt relatively immersed, did not feel nauseous, and were able to use the baseline video as a period of relaxation.

Therefore, for each video, we produced five mean memorability scores: a free recall score (from stage 1 of the debrief), and unrecalled, somewhat recalled, mis-recalled and correctly recalled scores (from the cued recall test in stage 2 of the debrief). We then considered how these scores correlated, to establish which variables should be considered in analyses for this paper. Free recall scores negatively correlated with unrecalled (r= −.292, p=.044) and somewhat recalled scores (r= −.449, p=.001), positively correlated with correct recall scores (r=.434, p=.002), and did not correlate with mis-recalled scores. As unrecalled and correctly recalled spaces had a very strong negative correlation (r= −.742, p=.000), and as the same was true for somewhat recalled and correctly recalled spaces (r= − .936, p=.000), we chose to consider free recall scores (from stage 1) and correctly recalled scores (from stage 2; known in this paper as cued recall scores) as the memorability metric for our analyses.

#### MRI pre-processing and analysis

Quality checks were performed using MRIQC 22.0.6; six participants were removed at this stage as they did not meet one of six key image quality metrics (AQI, FD>.2, FD_mean>.2, AOR, GCOR, dvars_nstd). One was further removed due to scanner error. Pre-processing was performed on the remaining 26 participants’ data (12 PG1, 14 PG2) using fMRIprep 22.1.1 with default settings^114^. First and second level GLM analyses were undertaken using SPM12. The primary categorical GLM included the following regressors: button presses, onsets of baseline video, and onsets of novel videos. Categorical parametric modulators were included on novel movies for: positive valence, negative valence, simple layout, complex layout, GB space, presence of people, task focus, and luminance (non-categorical, mean values). This model was used to carry out categorical contrast analyses: complex vs simple spaces, and positive vs negative spaces. Non-categorical parametric GLMs were also constructed to incorporate group mean scores for video valence, complexity, fascination, coherence, hominess, unusualness, arousal, curvature, luminance and memorability. This model was used to consider how the BOLD response covaries with the strength of a video’s valence, for example, rather than just the binary measure of whether it is a positive or negative. All mean values were demeaned for modelling.

For each contrast tested, small volume correction was run over whole-brain unthresholded maps (p<.001, k>5) using relevant ROIs. For contrasts pertaining to valence, ROIs included ACC, OFC, vmPFC, amygdala and insula. For contrasts concerning complexity, ROIs included hippocampus, parahippocampal gyrus, OPA and RSC. Memorability was evaluated with hippocampal, parahippocampal and amygdala ROIs. Curvature was evaluated with the aforementioned valence and complexity ROIs. All ROIs were anatomically defined using AAL 3 in the WFU Pick Atlas toolbox, except for the OPA, which was functionally derived from individual and group-level masks by Julian, Fedorenko, Webster, & Kanwisher^75^. We report peak level activations from small volume corrections and cluster size in tables. In more exploratory analyses, each contrast was also tested under whole-brain family-wise error (FWE) correction (p<.05, k>5). Without a priori hypotheses we do not discuss uncorrected results, but results tables reporting both peak-level activations significant at p<.05 FWE-corrected and p<.001 uncorrected are available upon request for full disclosure of the data. Tables were created using BSPMVIEW toolbox v.20161108 with AAL 3 atlas^115^, and regions verified using the MNI2TAL Tool^116,117^.

## Acknowledgments

We would like to thank Roger Atkins for MRI operator support; Jeremy Skipper and Chris Gahnstrom for continued advice; Spiers Lab for help with data collection; Greg Cooper for initial help in script development; and Delaram Farzanfar for performing the MLV Toolbox analysis to provide curvature scores.

## Data Availability

Videos can be found at https://osf.io/kbdvg/?view_only=7001f5a0d12b444f8e3586c5aa897ccc. Other materials are available upon request.

